# Alzheimer’s disease and its co-pathologies: implications for hippocampal degeneration, cognitive decline, and the role of *APOE* ε4

**DOI:** 10.1101/2025.04.11.648343

**Authors:** Klara Gawor, Sam Verrept, Geethika Arekatla, David Wouters, Alicja Ronisz, Moritz Hecht, Celeste Laureyssen, Helena Ver Donck, Bas Lahaije, Simona Ospitalieri, Mathieu Vandenbulcke, Markus Otto, Christine A.F. von Arnim, Estifanos Ghebremedhin, Bernard Hanseeuw, Rik Vandenberghe, Matthew Blaschko, Alejandro Sifrim, Kristel Sleegers, Dietmar Rudolf Thal, Sandra O. Tomé

**Affiliations:** Laboratory of Neuropathology, Department of Imaging and Pathology, KU Leuven, Leuven, Belgium; Laboratory of Neurobiology, Department of Neuroscience, KU Leuven, Leuven, Belgium; Laboratory of Multi-omic Integrative Bioinformatics, Department of Human Genetics, KU Leuven, Leuven, Belgium; Laboratory of Neuropathology, Institute of Pathology, Ulm University, Ulm, Germany; Complex Genetics of Alzheimer’s Disease Group, VIB Center for Molecular Neurology, VIB, Antwerp, Belgium; Department of Biomedical Sciences, University of Antwerp, Antwerp, Belgium; Laboratory for Translational Neuropsychiatry, Department of Neuroscience, KU Leuven, Leuven, Belgium; Department of Neurology, Ulm University, Ulm, Germany; Department of Geriatrics, University Medical Center Gottingen, Gottingen, Germany; Institute of Anatomy, Johann Wolfgang Goethe University, Frankfurt, Germany; Institute of Neuroscience, UCLouvain, Brussels, Belgium; Department of Neurology, UZ Leuven, Leuven, Belgium; Laboratory for Cognitive Neurology, Department of Neurosciences, KU Leuven, Leuven, Belgium; Processing Speech and Images, Department of Electrical Engineering, KU Leuven, Leuven, 3000, Belgium; Department of Pathology, UZ Leuven, Leuven, Belgium

**Keywords:** mixed dementia, hippocampal degeneration, digital pathology, α-synuclein, limbic-predominant age-related TDP-43 encephalopathy neuropathological change (LATE-NC), amyloid β (Aβ), Tau, small vessels disease, cerebral amyloid angiopathy (CAA), medial temporal lobe, *APOE* ε4

## Abstract

**INTRODUCTION:** In neurodegenerative dementias, the co-occurrence and interaction of Aβ, tau, and other pathological lesions confound their individual contributions to neurodegeneration and their modulation by risk factors.

**METHODS:** We analyzed 480 post-mortem human brains (ages 50–99) using regression and structural equation models to assess the relationships among Aβ, tau, LATE-NC, α-synuclein, other age-related lesions, and *APOE* ε4, as well as their effects on CA1 neuronal density, brain weight, and cognitive status.

**RESULTS:** Aβ, tau, LATE-NC, and amygdala-predominant α-synuclein pathology were highly interconnected. Tau was the strongest predictor of global neurodegeneration, while LATE-NC primarily, but not exclusively, affected hippocampal neuron loss. Small vessel disease correlated with both LATE-NC and α-synuclein, while *APOE* ε4 was mainly associated with extracellular and capillary Aβ.

**DISCUSSION:** Although Alzheimer’s pathology plays a central role in brain degeneration, coexisting pathologies can both exacerbate and independently contribute to it. These factors should be considered in patient stratification.

## BACKGROUND

Amyloid-β (Aβ) and hyperphosphorylated tau (pTau) deposits are strongly correlated with cognitive impairment in elderly patients.^1–3^ However, emerging evidence suggests that these hallmark Alzheimer’s disease neuropathological changes (ADNC) may not fully account for the extent of neuronal loss observed in these individuals. A range of additional alterations, including other protein aggregates and vascular lesions, frequently coexists with ADNC and may contribute to brain degeneration through synergistic or additive mechanisms.^4–6^

The medial temporal lobe (MTL) is particularly vulnerable to age-related changes. It includes key structures such as the parahippocampal region, hippocampal formation, and amygdala, which are essential for declarative memory formation, affective learning, and more.^7^ The MTL is among the first regions affected by pTau aggregation^2^ and by atrophy associated with neurodegenerative dementia.^8^ It is also the first region where other pathological lesions accumulate, including specific variants of TDP-43 and α-synuclein (αSyn) pathologies.^9–13^

TDP-43 aggregates that primarily accumulate in the MTL are classified as limbic-predominant age-related TDP-43 encephalopathy neuropathological changes (LATE-NC), a condition frequently observed alongside ADNC and distinct from other TDP-43 proteinopathies, such as amyotrophic lateral sclerosis or frontotemporal lobar degeneration.^9,11^ Similarly, while αSyn pathology in Lewy body disease typically follows a caudo-rostral progression (αSyn CR), with early involvement of the brainstem, individuals with severe ADNC often exhibit an amygdala-predominant pattern (αSyn AmyP), in which αSyn aggregates accumulate early in the MTL.^10,12^ Although the role of Aβ in promoting tau aggregation and spread has been recognized for years,^14^ growing evidence suggests that pTau, TDP-43, and αSyn often colocalize within the same lesions and may interact to exacerbate pathology.^15–19^

Hippocampal neuronal loss has been associated with multiple brain pathologies. Strong evidence links it to LATE-NC,^9,11^ and our recent observations indicate that αSyn AmyP correlates with reduced neuronal density in the hippocampus.^16^ Additionally, ADNC, including pTau^20^ and Aβ,^21^ as well as other lesions, have been implicated in hippocampal damage. These include argyrophilic grain disease (AGD),^22^ aging-related tau astrogliopathy (ARTAG),^23^ granulovacuolar degeneration (GVD),^24^ Hirano bodies,^25^ and cerebrovascular factors like cerebral amyloid angiopathy (CAA),^26^ cerebral small vessel disease (SVD),^23^ hippocampal calcification,^27^ and atherosclerosis of the circle of Willis (AS).^28^ However, given the frequent co-occurrence of these lesions, their individual contributions to neuronal loss remain uncertain.

Apolipoprotein E (APOE) is a major carrier of cholesterol and other lipids in the central nervous system.^29^ The *APOE* ε4 allele is one of the most extensively documented risk factors for Alzheimer’s disease (AD)^30^ and has been associated with numerous age-related neuropathological processes.^31–33^ While there is strong evidence connecting the *APOE* ε4 allele to the accumulation of extracellular Aβ deposits,^34^ its role in other neuropathological alterations remains less well understood.

This study aimed to investigate how age-related pathologies associated with AD contribute to brain degeneration, particularly in the hippocampus, while accounting for their interdependencies and determining the role of the *APOE* ε4 allele in this process. To achieve this, we trained a machine learning algorithm to detect pyramidal neurons, enabling a standardized quantification of neuronal density in the CA1 hippocampal subfield. We then applied linear regression analyses and structural equation modeling to examine the interactions among various MTL pathologies and assess their relative contributions to neuronal loss in CA1, overall brain weight, and cognitive impairment.

## METHODS

### Human autopsy cohort

This study included a total of 480 human autopsy cases from individuals aged 50 years or older. The cases were part of the ARCK/LEUCALS (Alzheimer Research Center KU Leuven/Leuven Center for Amyotrophic Lateral Sclerosis) cohort. They were obtained from university or municipal hospitals in Leuven (Belgium; Ethics Committee Research (EC) UZ/KU Leuven identifiers: S52791, S55312, S59292, S64363), Brussels (Belgium; UCL-2020-355), Bonn, Offenbach am Main, and Ulm (Germany; EC UZ/KU Leuven identifier: S59295, Ulm identifier: 58/08), as well as from GE Healthcare (United Kingdom; ClinicalTrials.gov identifiers NCT01165554 and NCT02090855). Brain samples were collected in compliance with local and federal regulations governing the use of human tissue for research in Belgium, Germany, and the United Kingdom. All experiments were conducted following ethical clearance from the UZ/KU Leuven EC (identifier: S65147).

To minimize potential confounders, we excluded cases with neuropathologically confirmed diagnoses of amyotrophic lateral sclerosis, frontotemporal lobar degeneration, Huntington’s disease, multiple system atrophy, multiple sclerosis, brain tumors, encephalitis, severe hypoxic or hypoglycemic brain damage, or epilepsy-induced hippocampal sclerosis.

For 331 cases, the Global Clinical Dementia Rating (CDR) score^35^ was retrospectively assigned based on standardized clinical examination reports from different clinicians, conducted 1 to 4 weeks before death. These reports included assessments of cognitive function and documented abilities such as self-care, dressing, eating habits, bladder and bowel continence, speech, writing, reading, memory problems, and spatial orientation. For 64 cases, Mini-Mental State Examination (MMSE)^36^ scores were available and were converted to CDR scores using established cut-off values.^37^

### Gross examination

During the gross examination, brains were weighed, and brain slices were inspected for macroscopic hemorrhagic and ischemic infarcts. Additionally, the severity of AS plaques in the circle of Willis was graded using published criteria.^38^ Values were excluded for this variable if fewer than eight of the twelve circle of Willis arteries were assessed.

### Immunohistochemistry

After gross examination, either the left (n = 434) or the right hemisphere (n = 46) was formalin-fixed. Tissue samples from various brain regions were used in this study, including the anterior MTL, the posterior MTL, the middle frontal gyrus, the occipital cortex, the midbrain, the pons, the medulla oblongata, the cerebellum, and the basal ganglia. The samples were then embedded in paraffin and sectioned into 5 μm-thick slices for analysis.

The sections were stained using a panel of antibodies (see Supplementary Table 1 for the full list of stained regions, antibodies, and their concentrations). To prepare the sections, they were deparaffinized and subjected to epitope retrieval using citrate buffer (pH 6; Envision™ Flex Target Retrieval Solution, Dako, K8005) for 10 minutes. For αSyn and Aβ staining, an additional step was performed, involving a 5-minute incubation with formic acid. Tissue sections were incubated overnight with the primary antibody in a humid chamber. The following day, anti-mouse horseradish peroxidase (HRP)-linked secondary antibodies or the VectaStain ABC-HRP kit (Vector Laboratories) were applied. The binding between the primary and secondary antibodies was visualized using a DAB solution (Liquid DAB + Substrate Chromogen System, Dako). Finally, all slides were counterstained with hematoxylin using an autostainer, and coverslips were mounted automatically with a cover-slipper (Leica Microsystems). Additionally, the autostainer was used to prepare hematoxylin-eosin (HE) sections for all slides.

### Neuropathological assessment

Tissue examination and digital photography were conducted using a ZEISS Axio Imager 2 microscope with an Axiocam 506 camera and a DM2000 LED Leica microscope with a Leica DFC7000 T digital camera. Each case underwent a pathological assessment and diagnosis, with all parameters and their detailed operational definitions provided in Supplementary Table 2 and representative lesion images shown in Figure 1.

**FIGURE 1.**
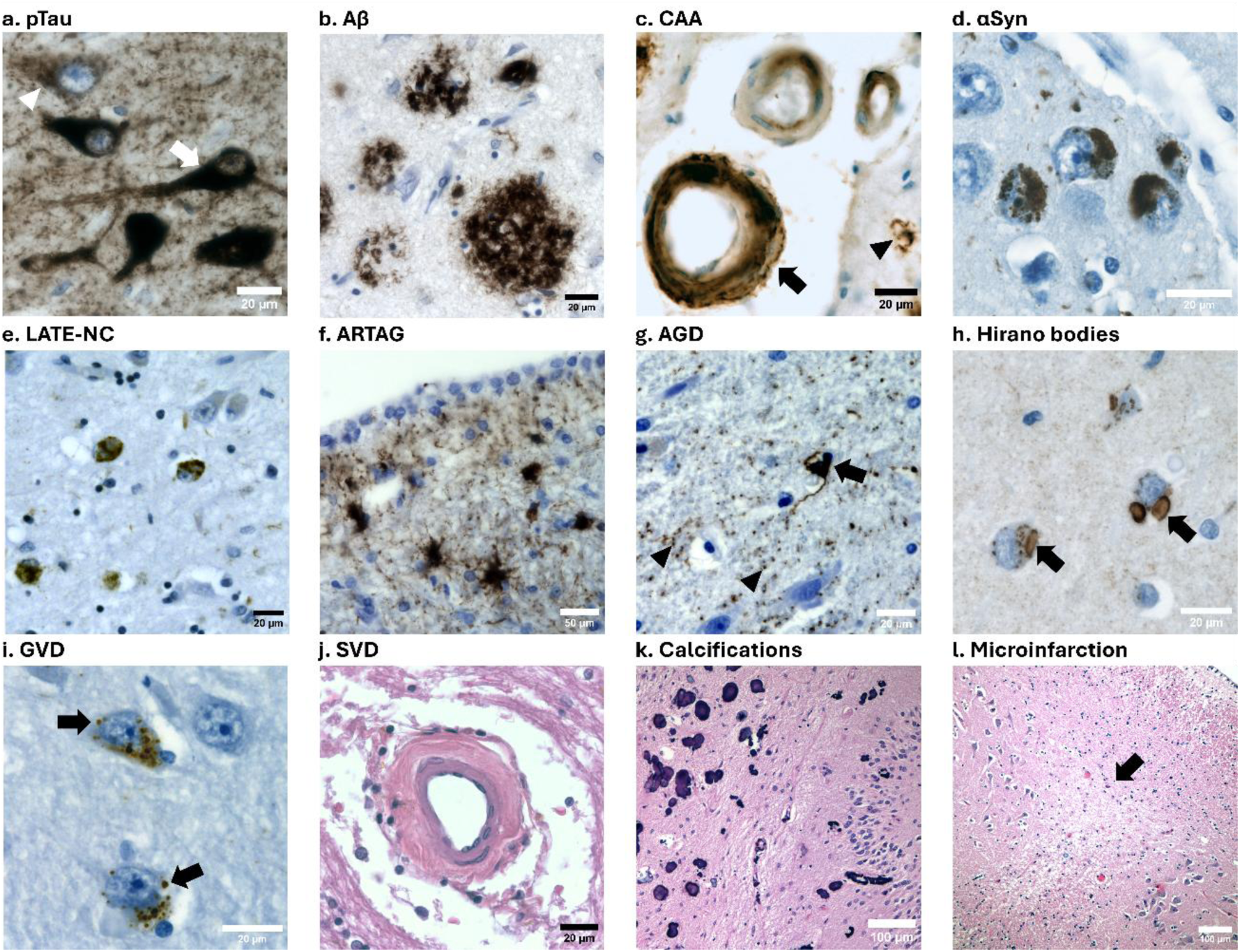
Neuropathological lesions present in the MTL. **(A)** NFTs (arrows), pre-tangles (arrowhead), and dystrophic neurites positive for pTau in CA1 neurons. Antibody: pTau [S202/T205]. ×400. **(B)** Aβ plaques in the entorhinal cortex. Antibody: Aβ [17–24]. ×400. **(C)** CAA affecting arterioles (arrow) and capillaries (arrowhead) in the occipital cortex. Similar lesions are also present in the MTL. Antibody: Aβ [17–24]. ×400. **(D)** αSyn aggregates in amygdala neurons. Antibody: anti-aggregated αSyn. ×400. **(E)** pTDP-43 neuronal cytoplasmic inclusions in amygdala neurons from a case following the LATE-NC pattern. Antibody: pTDP-43 [409/410]. ×200. **(F)** Subventricular ARTAG in the anterior MTL. Antibody: pTau [S202/T205]. ×400. **(G)** AGD with coiled bodies (arrow) and grains (arrowhead) in CA1. Antibody: pTau [S202/T205]. ×400. **(H)** Hirano body inclusions (arrows) in CA1. Antibody: F-actin. ×400. **(I)** Granulovacuolar degeneration bodies (arrows) in CA1 neurons. Antibody: pTDP-43 [409/410]. ×400. **(J)** An arteriole in the parahippocampal white matter affected by arteriolosclerosis, a form of SVD characterized by hyalinization, thickening, and loss of mural cells in the vessel wall. HE staining. ×200. **(K)** Calcified vessels in the molecular layer of CA1 and DG regions of the posterior hippocampus. HE staining. ×100. **(L)** Old microinfarct (arrow) located at the border of CA2 and CA1. HE staining. ×100.

Phases of Aβ plaque deposition, Braak stages for neurofibrillary tangles (NFTs) and CERAD score for neuritic plaques (NPs) were obtained following established protocols.^2,3,39^ The neuropathological diagnosis of ADNC was performed using the criteria published by the National Institute on Aging and Alzheimer’s Association, with the ABC score applied as a general measure of ADNC severity.^1^ CAA severity was assessed according to Vonsattel grades and subtyped as type 1 (with capillary involvement) or type 2 (without capillary involvement).^40^

The presence of AGD and ARTAG in the MTL was assessed using AT8 staining.^41,42^ LATE-NC was diagnosed and staged based on recently published consensus guidelines.^9,43^ The presence of pTDP-43 lesions in the dentate gyrus (DG) of the posterior hippocampus was also evaluated. Additionally, pTDP-43 staining was used to stage GVD progression according to established criteria.^44^

The severity of αSyn pathology in the medulla oblongata, midbrain, amygdala, and temporal cortex was semi-quantitatively assessed and converted into a MTL-to-brainstem severity ratio, as described previously.^16^ This ratio was used to distinguish between two spreading variants: αSyn AmyP (ratio > 1) and αSyn CR (ratio ≤ 1).

The severity of SVD in the white matter of the temporal cortex was semi-quantitatively scored on HE-stained sections from the posterior and anterior MTL, based on the number of affected vessels, as detailed earlier.^16^ Microinfarctions and lacunar infarctions were identified on HE-stained sections as part of the diagnostic process. Using posterior MTL HE section we evaluated the presence of calcified vessels in the molecular layer of DG and CA1. Finally, we assessed the severity of Hirano body pathology in the CA1 region using categories based on lesion counts in three photographs taken at 200x magnification (623 × 468 µm each): 0 (no lesions), mild (1–8 lesions), moderate (9–25 lesions), and severe (≥26 lesions).

### *APOE* genotyping

Genomic DNA was extracted from either fresh-frozen or paraffin-embedded brain tissue. For 144 cases, genotyping of the *APOE* single nucleotide polymorphisms (SNPs) rs429358 and rs7412 was performed using low-coverage whole-genome sequencing (WGS), as previously described.^33^ Library preparation was conducted using the xGen cfDNA & FFPE DNA Library Preparation Kit (IDT, Belgium) with barcoding, followed by equimolar pooling of the samples and sequencing with Avidity Chemistry (Arslan, 2023) at an average depth of 1x coverage on an AVITI instrument (Element Bioscience, San Diego). Raw FASTQ files were aligned to the GRCh38 genome build using the Burrows-Wheeler Alignment (BWA) tool and imputed using the Genotype Likelihoods IMputation and PHasing mEthod (GLIMPSE)^45^ against the 1000 Genomes haplotype reference panel. Allele dosages for both *APOE* SNPs were extracted to derive the *APOE* ε2, ε3, and ε4 alleles. All SNPs had genotype probability (GP) higher than 0.9.

For 18 samples, *APOE* genotyping was performed using Sanger sequencing. DNA was amplified using simplex polymerase chain reaction (PCR; forward primer: GCCTACAAATCGGAACTGGA, reverse primer: CTCGAACCAGCTCTTGAGGC) with Platinum Taq (Thermo Fisher Scientific, Waltham, MA, USA). Following purification of the PCR products with Illustra™ ExoProStar™ (Cytiva, Marlborough, MA, USA), dideoxy chain termination was conducted with the BigDye Terminator v3.1 Cycle Sequencing Kit (Thermo Fisher Scientific). Sequencing was carried out on an ABI 3730XL DNA Analyzer (Thermo Fisher Scientific) according to the manufacturer’s protocol. Sequence analysis was performed using novoSNP v3.9.1 and SnapGene v6.03 software (GSL Biotech LLC, San Diego, CA, USA).

Additional *APOE* genotyping data were available for 120 cases, obtained using PCR followed by *HhaI* restriction enzyme digestion, as previously described.^46^ For another 85 cases, *APOE* genotype information was provided by GE Healthcare in the context of clinical trials NCT01165554 and NCT02090855.

### Automated neuronal detection algorithm

#### Input dataset

Pyramidal neurons in the CA1 region were manually annotated using the annotation tool in QuPath (v0.5.1) across 801 images derived from pTDP-43 409/410-stained sections of 267 individuals, in accordance with the criteria described in.^16^ In addition to neuronal annotation, non-neuronal structures, such as blood vessels, were annotated in 72 pTDP-43-stained images of the CA1 region and 16 images from the molecular layer of the DG and CA1. Each image covered an area of 500 × 500 μm and was captured using a ZEISS Axio Imager 2 Microscope equipped with an Axiocam 506 camera, a 20x objective, and auto-exposure settings. The dataset encompassed a range of conditions, including varying levels of neuronal loss, differences in fixation times, durations of paraffin embedding, post-mortem intervals, and phases of photobleaching.

#### Object detection

Images were first converted to grayscale format and inverted when the objects of interest appeared as black on a white background instead of the expected white on black. To reduce artifacts and noise, a small Gaussian blur was applied. Foreground-background segmentation was performed by computing a threshold based on a subsampled version of the image using the threshold_local function from scikit-image (version 0.24.0). Local maxima were identified using skimage.feature.peak_local_max, and those belonging to the same object were merged using the DBSCAN clustering algorithm from scikit-learn (version 1.5.2). The centroid of each merged cluster was determined as the midpoint of its contributing local maxima, and a unique label was assigned to each identified object.

#### Creation of classifier training data

Extracting objects for the creation of the training data was slightly adapted from the protocol above. For this purpose a wider net was cast in detecting objects to generate enough negative controls for training. Specifically, the above protocol is cut short after thresholding foreground and background. Small holes are filled using scipy.ndimage.binary_fill_holes. Separate connected regions of foreground-labeled pixels are then given a unique label, indicating the presence of an object.

Extracting single-object images was done by matching the manually indicated center coordinates with the corresponding label of the thresholded object. A ‘positive’ image was created using a 100 × 100 pixel bounding box around the center coordinate, after which all labeled pixels in this bounding box and their connected pixels were set to zero to avoid matching any nearby objects in another extraction. Negative images were created using the same bounding box extraction applied to a random selection of remaining objects or manually annotated non-neurons. A total of 122,801 single-object images were extracted, with 31,273 annotated as ‘Positive’ and 91,528 as ‘Negative.’ These were split into training (60%), validation (20%), and test (20%) subsets. All training images were saved as 32-bit depth RGB png files.

#### Training of image classifier

A binary classifier was trained using transfer learning with ResNet-34 as the base architecture to distinguish photos of neurons from non-neurons. ResNet-34, a pre-trained convolutional neural network trained on ImageNet, was fine-tuned for this task. Model training was performed using PyTorch via FastAI v2. Training was conducted over three epochs using the vision_learner and fine_tune functions. During the first epoch, only the final layer was trained while the remaining layers were frozen. In the subsequent epochs, all layers were unfrozen and trained. A default batch size of 64 and a base learning rate of 0.002 were used. The validation error rate was 10.8% after the first epoch, decreasing to 5.5% by the third epoch, after which it plateaued. Each training round took approximately 30 minutes on a local CPU with six cores and 16 GB of RAM.

#### Estimating neuronal density

Three images of the posterior CA1 region were collected from each of the 480 cases included in this study, following the same guidelines as those used for the input dataset. Object detection was performed, and the identified objects were classified as either neurons or non-neurons using the trained classifier. Objects within 10 μm of the image edge were excluded. If two detected neurons were within 15 μm of each other, one was randomly removed to prevent duplicate counting. Finally, neuron density was calculated for each case in neurons per mm².

#### Validation on independent dataset

Additionally, sixty randomly selected photos, not included in the training dataset, were manually annotated by an independent rater who was not involved in the training annotation process. The number of neurons identified by the automated neuronal detection algorithm in each photo was compared and correlated with the counts identified by the human rater.

### Statistical analysis

Statistical analyses were conducted in RStudio (version 2024.09.0) using R 4.3.0. Differences in brain degeneration across pathology combinations were assessed using Kruskal-Wallis tests, followed by pairwise Wilcoxon rank-sum tests with Bonferroni-Holm (BH) correction. Spearman’s correlation was used to examine the relationships between demographic variables (age at death and sex) and neuropathological parameters. Additionally, semi-partial Spearman’s correlation, controlling for age at death, was performed to assess associations between each pair of neuropathological parameters. All correlation p-values were adjusted using the BH method. Furthermore, three series of linear regression analyses were conducted to examine the relationships between brain degeneration measures (including CA1 neuronal density, brain weight, and CDR) and neuropathological parameters, each incorporating a different set of covariates. All variables were standardized, and p-values for neuropathological parameters were BH-adjusted within each regression series.

Additionally, structural equation model analyses were performed using the lavaan package (version 0.6.16). The directionality of arrows in models was decided based on the literature on protein aggregate interactions (Supplementary Table 3), however, the path αSyn → LATE had to be reversed to avoid non-recursive models. Model parameters were estimated using maximum likelihood estimation with robust standard errors, a Satorra-Bentler scaled test statistic, and the NLMINB optimization method. Model fit was assessed using the robust Comparative Fit Index (CFI; good fit > 0.97), robust Standardized Root Mean Square Residual (SRMR; good fit < 0.05), and standardized Root Mean Square Error of Approximation (RMSEA; good fit < 0.05).

## Data availability

The raw data used in this study are publicly available in Supplementary Table 18.

## RESULTS

The study cohort included 480 individuals aged 50 to 99 years, with 52.1% being male and 50.6% were classified as cognitively impaired (CDR > 0, assessed in 395 cases). A detailed summary of demographics, neuropathological parameters, and the number of cases analyzed is provided in Table 1. The operationalization of the parameters is detailed in Supplementary Table 2.

**TABLE 1.** Summary statistics of cohort demographics and neuropathological parameters.

| Parameter | Mean (SD) | Frequency (%) | Min-Max | n |
| --- | --- | --- | --- | --- |
| Age at death (y) | 74.6 (10.8) | - | 50-99 | 480 |
| Sex (male) | - | 52.1% | - | 480 |
| CDR | 1.06 (1.26) | - | 0-3 | 395 |
| Cognitively intact ($\text{CDR} = 0$) | - | 49.4% | - | - |
| MCI ($\text{CDR} = 0.5$) | - | 6.3% | - | - |
| Dementia ($\text{CDR} > 0.5$) | - | 44.3% | - | - |
| Neuronal density CA1 (per $\text{mm}^2$) | 163.3 (67.4) | - | 1.3-387.5 | 480 |
| Brain weight (g) | 1238.5 (171.6) | - | 585-1700 | 333 |
| PMI (hours) | 35.2 (30.5) | - | 1.5-144 | 423 |
| *APOE* | - | - | - | 367 |
| 2/2 | - | 0.8% | - | - |
| 2/3 | - | 10.1% | - | - |
| 2/4 | - | 2.2% | - | - |
| 3/3 | - | 55% | - | - |
| 3/4 | - | 27% | - | - |
| 4/4 | - | 4.9% | - | - |
| Braak NFT stages (pTau) | 2.7 (1.9) | - | 0-6 | 480 |
| Aβ phases | 2.6 (2) | - | 0-5 | 480 |
| CERAD score (NPs) | 0.9 (1.2) | - | 0-3 | 480 |
| LATE-NC stages | 0.5 (0.9) | - | 0-3 | 480 |
| pTDP-43 DG | - | 10.2% | 0-1 | 479 |
| αSyn variant | - | - | - | 480 |
| No αSyn | - | 67.30% | - | - |
| Amygdala-predominant | - | 10.6% | - | - |
| Caudo-rostral | - | 22.1% | - | - |
| SVD in temporal cortex | 0.9 (0.9) | - | 0-3 | 366 |
| AS severity | 1.6 (0.8) | - | 0-3 | 298 |
| CAA type | - | - | - | 480 |
| No CAA | - | 44.0% | - | - |
| Type 1 | - | 30.0% | - | - |
| Type 2 | - | 26.0% | - | - |
| CAA severity | 1 (1) | - | 0-3 | 480 |
| Hippocampal calcifications | - | 23.5% | 0-1 | 442 |
| Infarctions | - | 18.4% | 0-1 | 418 |
| ARTAG | - | 19.4% | 0-1 | 324 |
| AGD | - | 12.6% | 0-1 | 445 |
| Hirano bodies severity | 1 (0.7) | - | 0-3 | 278 |
| GVD stages | 1.4 (1.7) | - | 0-5 | 355 |

Note: Continuous and ordinal variables are presented as mean and standard deviation (SD).
Binary and categorical variables are reported as the percentage of cases exhibiting the specified characteristics. MCI – mild cognitive impairment

### Neuronal detection algorithm

To assess the density of pyramidal neurons in the posterior CA1 subfield of the hippocampus across all cases, a pyramidal neuron detection algorithm using a neural network classifier was trained (Figure 2A). Examples of neuron detections are shown in Figure 2B and Supplementary Fig. 1. Classifier metrics for the validation dataset were as follows: Accuracy: 94.4%, Sensitivity: 88.4%, Specificity: 96.5%, Precision: 90%. Similar metrics were also obtained on the test dataset. For 60 randomly selected images, independently evaluated by a reviewer not involved in classifier training, the Spearman’s correlation coefficient between the algorithm’s output and the human ratings was 0.83 (Supplementary Fig. 2), indicating high reliability of the detection algorithm.

**FIGURE 2.**
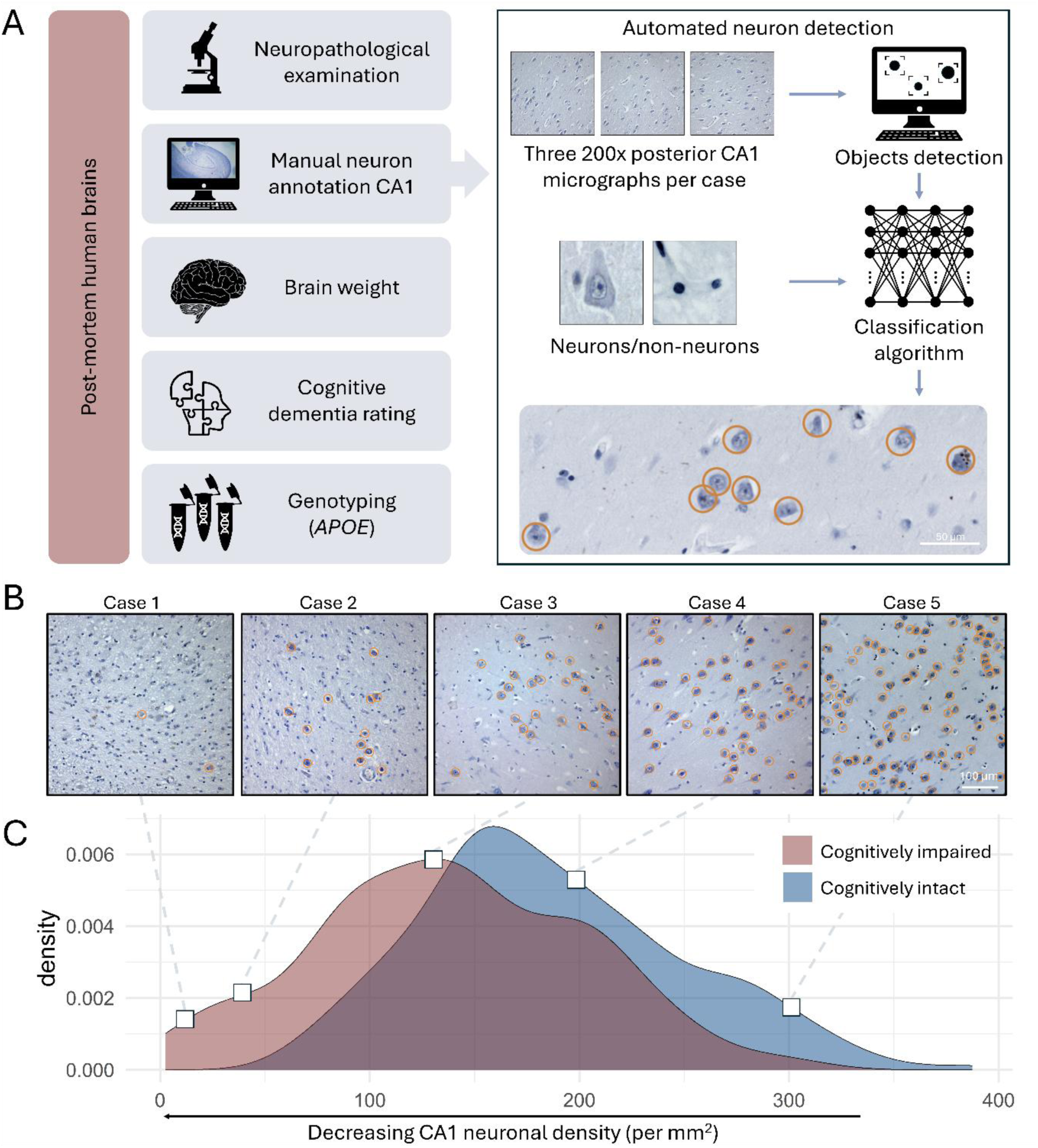
(**A**) Schematic representation of the study design. **(B)** Representative images from the CA1 subfield of the posterior hippocampus in five different cases, with neurons identified by the trained classifier marked by orange circles. **(C)** Density plot showing the distribution of CA1 neuronal densities in cognitively impaired (CDR > 0, n = 195) and cognitively intact (CDR = 0, n = 200) cases. Cognitive impairment status was unavailable for 85 cases.

The mean neuronal density in the CA1 subfield in our sample was 163.3 neurons per mm² (SD = 67.4), with values ranging from 1.3 to 387.5. Figure 2C illustrates the distribution of neuronal damage in cognitively intact and cognitively impaired individuals. The distribution of brain weights, stratified by cognitive status, is presented in Supplementary Fig. 3.

### Prevalence of mixed pathologies

The sample encompassed a broad range of ADNC and LATE-NC severities, as well as two αSyn pathology variants (Figure 3A). Within the cohort, 93.1% of cases exhibited pTau aggregates, 75% had Aβ deposits, 32.7% displayed αSyn pathology, and 26.3% had LATE-NC.

**FIGURE 3.**
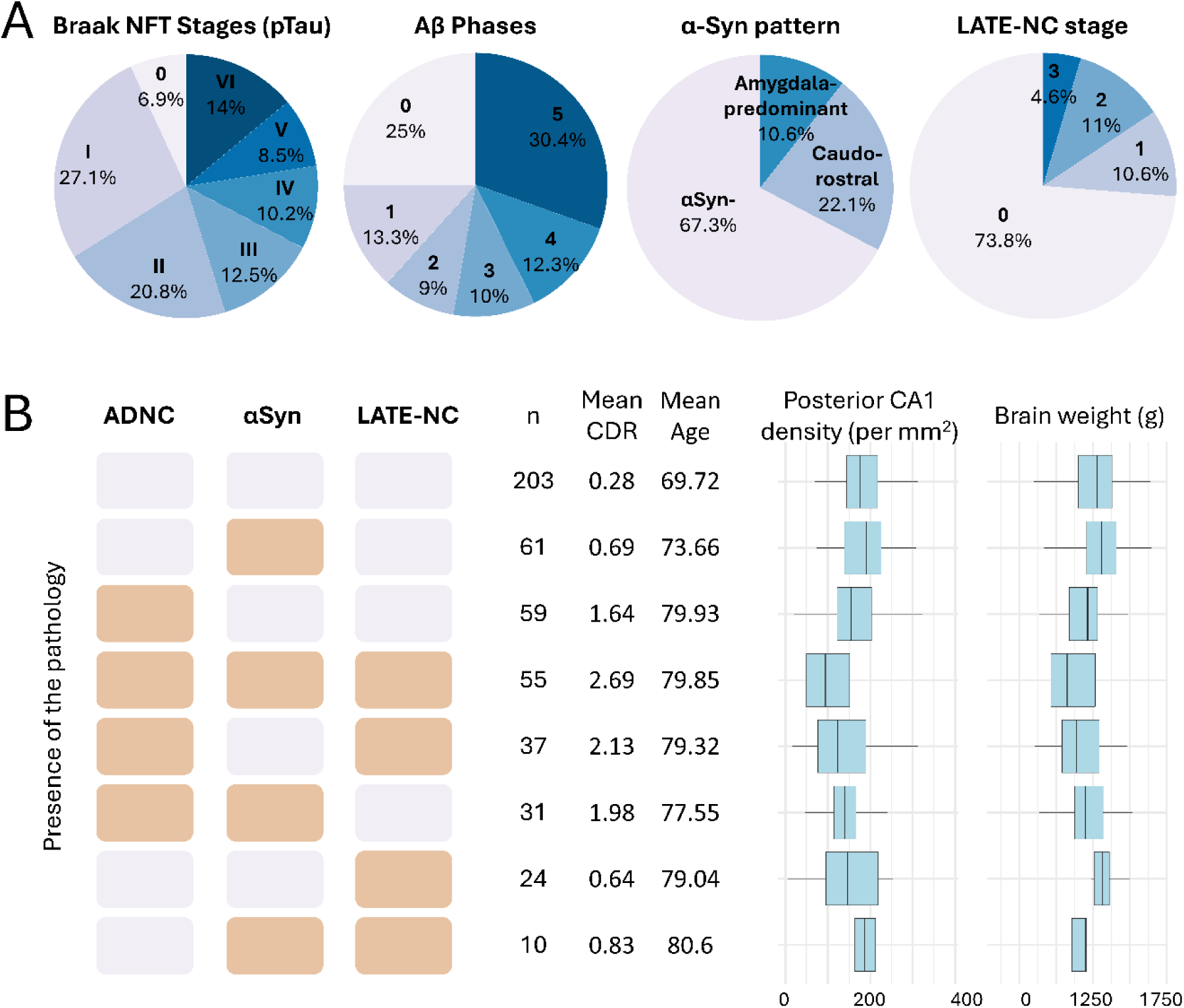
Combinations of neuropathological lesions and their brain degeneration correlates. **(A)** Proportion of cases with specific pathological lesions identified in this study in total sample of 480 cases. **(B)** Overview of combinations of moderate or severe ADNC, αSyn pathology, and LATE-NC in the study cohort, arranged in descending order of prevalence (left panel). The number of cases in each group, their mean CDR score, and mean age at death are reported (middle panel). Boxplots illustrate neuronal density in the CA1 and brain weight for each combination (right panel). Horizontal bold lines in the boxplots represent median values, box margins indicate the 25th and 75th percentiles, and whiskers display the full range of observed values. A summary of the Kruskal-Wallis analysis with pairwise Wilcoxon tests comparing brain degeneration measures across different combinations can be found in Supplementary table 5.

Mixed pathologies were common (Figure 3B, Supplementary Table 4). Among the 182 cases with moderate to severe ADNC, 92 (51%) also had LATE-NC. Additionally, 86 (47%) exhibited αSyn pathology, including 45 with the AmyP variant and 41 with the CR variant. A triple pathological combination—comprising ADNC, αSyn, and LATE-NC—was observed in 55 cases, accounting for 30% of all cases with moderate to severe ADNC.

Individuals with triple pathology had the lowest neuronal density in the CA1 region (mean = 108.1), significantly lower than individuals with ADNC only (mean = 160.9, *P* value < .001) or those without any of the three pathologies (mean = 182.2, *P* value < .001), as confirmed by a Kruskal-Wallis test followed by pairwise Wilcoxon comparisons with BH correction (Supplementary Table 5). Additionally, individuals with triple pathology showed the greatest cognitive impairment across all pathology combinations (all BH-corrected *P* value < .05).

### Correlations between age-related neuropathologies, age at death, and sex

We analyzed relationships between key age-related neuropathological markers, age at death, and sex using Spearman’s correlation (Supplementary Table 6). The neuropathological parameters included Aβ, pTau, NPs, LATE-NC stages, TDP-43 positivity in the DG, αSyn variants, temporal lobe SVD, AS, CAA severity, CAA type 1, hippocampal vascular calcification, multiple old brain infarctions, ARTAG, AGD, Hirano bodies, and GVD.

A positive correlation with age at death was observed for the majority of neuropathological parameters, except for αSyn CR (*P* value *=* .145), hippocampal calcification (*P* value *=* .897), Hirano bodies (*P* value *=* .669), and AGD (*P* value *=* .08). Regarding sex differences, the only significant finding was a greater prevalence of αSyn CR in males (*P* value *=* .012). Additionally, trends approaching significance (*P* value *<* .1) were observed for associations between females and increased pTau and GVD, as well as males and a higher prevalence of infarctions.

Next, we investigated associations between individual lesions using Spearman correlation analysis, adjusted for age (Figure 4). Correlation coefficients, exact p-values, and the number of pairwise cases are provided in Supplementary Table 7. The analysis revealed strong correlations among ADNC, CAA, GVD, αSyn AmyP, and LATE-NC parameters (BH-corrected *P* values *<* .05 for all correlation pairs). Additionally, temporal SVD correlated with ADNC, LATE-NC, αSyn AmyP, and ARTAG. Cerebrovascular parameters such as SVD, AS, and multiple old infarctions were intercorrelated. AGD showed a negative correlation with ADNC, CAA, and αSyn AmyP pathology.

**FIGURE 4.**
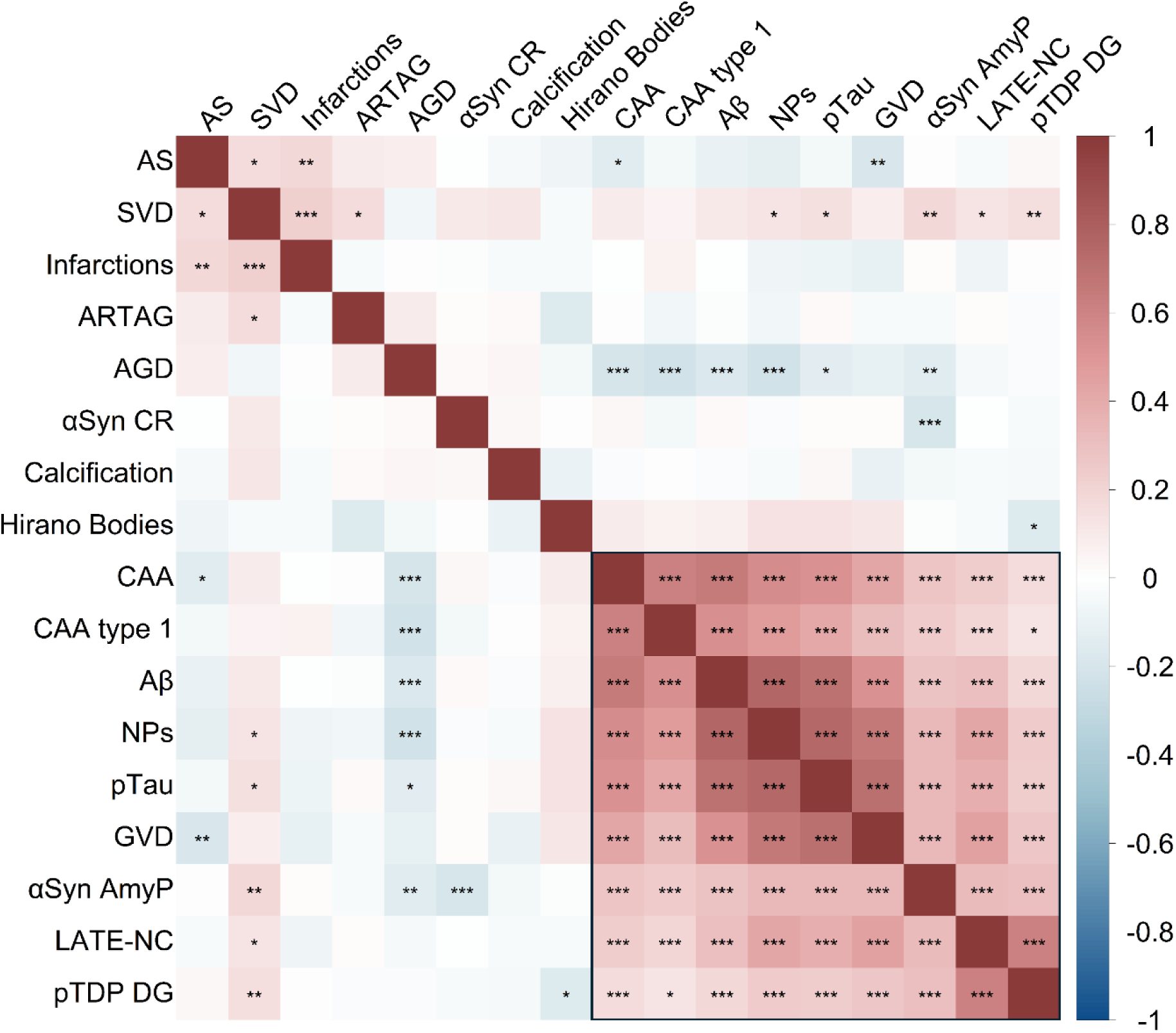
Heatmap displaying the results of a semi-partial Spearman’s correlation of neuropathological lesions, corrected for age at death. Stars indicate significance levels of p-values adjusted using the BH correction (* *P* value < .05, ** *P* value < .01, *** *P* value < .001). The shade of each tile represents the strength and direction of the correlation. The order of tiles has been arranged using hierarchical clustering with complete linkage. A square highlights the spectrum of AD-related lesions that are strongly intercorrelated.

### Role of brain pathologies in hippocampal and global brain degeneration

To assess the impact of neuropathological lesions on brain degeneration, we conducted three sets of multiple linear regression analyses. Neuropathological lesions served as independent variables, while brain degeneration was estimated using three dependent variables: posterior CA1 neuronal density (reflecting medial temporal lobe damage), brain weight (indicating overall atrophy), and CDR score (serving as a marker of cognitive impairment).

The first series of analyses was adjusted for age at death, sex, and cohort. The second series was additionally controlled for overall ADNC levels. In the third series, we further adjusted for the post-mortem interval (PMI); this analysis was restricted to non-ADNC variables that remained significant in the second set. Complete statistical results, including standardized regression coefficients and sample sizes, are available in Supplementary Tables 8–10 and summarized in Supplementary Fig. 4–5.

All ADNC parameters (pTau, Aβ, and NPs) were associated with all brain degeneration measures (Figure 5A, Series 1; *P* value *<* .001 for all), emphasizing the need to control for these factors when analyzing the role of other neuropathologies.

**FIGURE 5.**
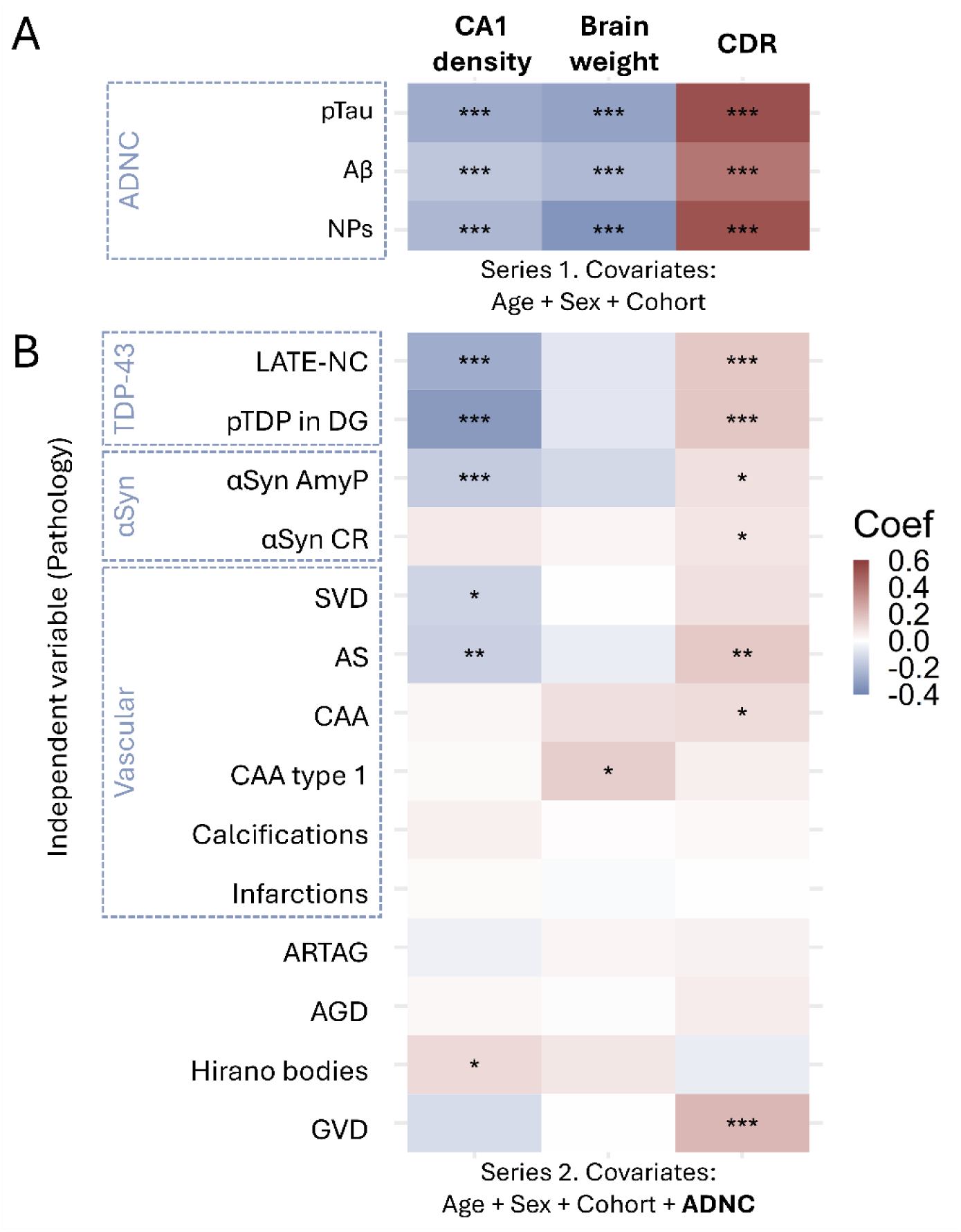
Heatmap showing standardized coefficients from multiple linear regression analyses evaluating brain degeneration measures as dependent variables and neuropathological lesions as independent variables. Regression models were adjusted for age at death, sex, and cohort (**A**, Series 1) and for the overall severity of ADNC (**B**, Series 2). The colors of the tiles represent the direction and strength of the coefficients for each neuropathological lesion. Asterisks indicate p-values after BH correction within each panel (* *P* value < .05, ** *P* value < .01, *** *P* value < .001).

After adjusting for ADNC (Figure 5B, Series 2), both LATE-NC stages and TDP-43 inclusions in the DG were negatively associated with CA1 neuronal density (β = –0.26, *P* value < .001; β = –0.32, *P* value < .001) and associated with increased cognitive impairment (β = 0.16, *P* value < .001 for both). αSyn AmyP pathology was linked to CA1 degeneration (β = –0.16, *P* value < .001) and cognitive impairment (β = 0.09, *P* value = .03). The αSyn CR variant was also associated with cognitive impairment (β = 0.08, *P* value = .04).

Moreover, vascular parameters such as SVD in the temporal cortex and AS severity were associated with CA1 neuronal loss (β = –0.13, *P* value = .02; β = –0.15, *P* value = .01). AS severity was also associated with greater cognitive impairment (β = 0.16, *P* value = .001). CAA severity was positively associated with cognitive impairment (β = 0.10, *P* value = .049), while CAA type 1 was correlated with increased brain weight (β = 0.14, *P* value = .03). GVD stages were related to cognitive impairment (β = 0.21, *P* value < .001). Interestingly, Hirano bodies were linked to increased CA1 neuronal density (β = 0.11, *P* value = .049). Other parameters, including hippocampal vascular calcifications, multiple old infarctions, ARTAG, and AGD, showed no association with brain degeneration measures in models with or without ADNC adjustment.

Models showing significant associations between pathological factors and brain degeneration measures after adjusting for ADNC were further controlled for PMI (Series 3, Supplementary Table 10), which resulted in a reduced sample size. After including this covariate, the associations between SVD and CA1 neuronal density (β = –0.1, *P* value = .052) and between αSyn CR and cognitive impairment (β = 0.07, *P* value = .057) were no longer statistically significant but remained at a trend level. All other associations remained significant.

### Modeling connections between AD-related lesions

Structural equation models examined the interactions among neuropathological parameters, their association with *APOE* ε4, and their effects on brain degeneration measures. Each model demonstrated a good fit based on standard criteria (Supplementary Table 11). Detailed summary statistics are available in Supplementary Tables 12–17. All reported path estimates are standardized (expressed in standard deviation units) to enable direct comparison of effect sizes across paths.

Strong Interconnectedness was observed among all core parameters (Model 1, Figure 6, n = 480), with the most robust associations found between Aβ and CAA severity (Estimate = 0.67, *P* value < .001) and Aβ and pTau (Estimate = 0.52, *P* value < .001), followed by LATE-NC and Aβ (Estimate = 0.29, *P* value < .001) and LATE-NC and αSyn AmyP (Estimate = 0.27, *P* value < .001). Additional associations were observed between LATE-NC and pTau (Estimate = 0.15, *P* value < .001), αSyn AmyP and CAA (Estimate = 0.15, *P* value = .002), pTau and αSyn AmyP (Estimate = 0.11, *P* value < .001), pTau and CAA (Estimate = 0.13, *P* value = .001), and between αSyn AmyP and Aβ (Estimate = 0.11, *P* value = .034). No association was found between LATE-NC and CAA (Estimate = 0.05, *P* value = .174). Subsequent models were developed based on the significant associations among neuropathological parameters identified in Model 1.

**FIGURE 6.**
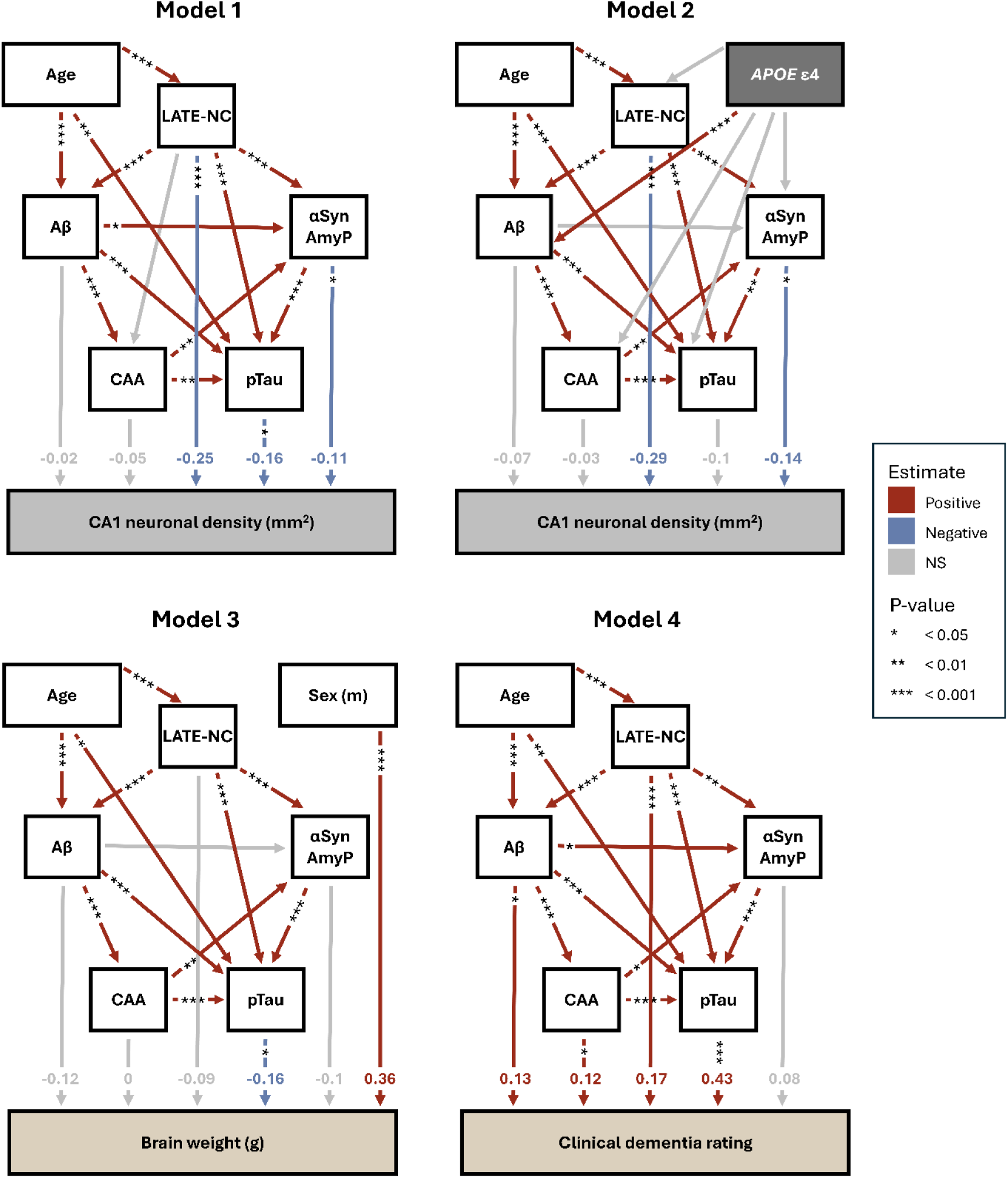
Structural equation models were used to investigate the relationships between neuropathological lesions associated with the AD spectrum and measures of brain degeneration. Model 1 (n = 480) examined the associations between AD-related lesions and neuronal density in the posterior CA1 region. The association between LATE-NC and CAA was not significant and was excluded from subsequent models. Model 2 (n = 367) explored the effect of the *APOE* ε4 allele on the accumulation of neuropathological lesions. A complementary model incorporating CAA type 1 is presented in Supplementary Fig. 6. Additionally, we analyzed the association between brain lesions and total brain weight in Model 3 (n = 333) and between neuropathological lesions and global cognitive impairment, as measured by the CDR, in Model 4 (n = 397). All directional relationships included in the models are represented by arrows. The color of each arrow indicates the sign of the estimate, except for non-significant (NS) path estimates (*P* value ≥ 0.5), which are shown in gray. Standardized estimates for paths involving brain degeneration parameters are indicated at the tip of each arrow.

After accounting for interrelationships among neuropathological parameters, Model 1 (Figure 6) also assessed their direct associations with CA1 neuronal density. LATE-NC (Estimate = – 0.26, *P* value < .001), pTau (Estimate = –0.16, *P* value = .014), and αSyn AmyP (Estimate = – 0.11, *P* value = .027) were associated with reduced CA1 neuronal density, whereas Aβ (Estimate = –0.02, *P* value = .81) and CAA severity (Estimate = –0.05, *P* value = .394) showed no effect. Additionally, a supplementary model was developed to assess whether SVD in the temporal lobe directly influences CA1 neuronal density (Model S1, Supplementary Fig. 6, n = 366). The results showed a nonsignificant direct association between SVD and CA1 neuronal loss (Estimate = –0.06, *P* value = .237). However, SVD was associated with both LATE-NC (Estimate = 0.14, *P* value = .013) and αSyn AmyP (Estimate = 0.11, *P* value = .023).

Including *APOE* ε4 status in the model (Model 2, Figure 6, n = 367) revealed a significant association with Aβ pathology (Estimate = 0.3, *P* value < .001). While the association between *APOE* ε4 and LATE-NC did not reach statistical significance, it approached the threshold (Estimate = 0.09, *P* value = .066). In contrast, we found no significant associations between *APOE* ε4 and pTau (Estimate = 0.03, *P* value = .391), αSyn AmyP (Estimate = 0.06, *P* value = .262), or CAA severity (Estimate = 0.06, *P* value = .131). However, when CAA severity was replaced with the presence of CAA Type 1 (Model S2, Supplementary Fig. 6, n = 367), a significant direct association with *APOE* ε4 emerged (Estimate = 0.24, *P* value < .001).

Next, we examined the pathological lesions associated with decreased brain weight (Model 3, Figure 6, n = 333). To account for a potential confounder, we included a path between brain weight and sex in this model. pTau was the only pathology with a direct effect that reached significance level (Estimate = –0.16, *P* value = .047). Although αSyn AmyP (Estimate = –0.10, *P* value = .071), LATE-NC (Estimate = –0.09, *P* value = .12), and Aβ (Estimate = –0.12, *P* value = .184) also had negative estimates for the path to brain weight, they did not reach statistical significance.

Finally, a model examining neuropathological pathways influencing cognitive impairment (Model 4, Figure 6, n = 395) identified a strong effect of pTau (Estimate = 0.43, *P* value < .001). Nevertheless, other pathologies also contributed to an increased likelihood of cognitive impairment, including LATE-NC (Estimate = 0.17, *P* value < .001), Aβ (Estimate = 0.13, *P* value = .04), and CAA severity (Estimate = 0.12, *P* value = .028). The association between αSyn AmyP and cognitive impairment was borderline significant (Estimate = 0.08, *P* value = .051).

## DISCUSSION

Our findings highlight that neurodegeneration in age-related dementias results from the complex interaction of multiple pathologies. We found that Aβ, pTau, LATE-NC, and amygdala-predominant αSyn contributed to brain degeneration both independently and through their interconnections. While pTau was the primary driver of global brain atrophy, hippocampal degeneration was predominantly, though not exclusively, associated with LATE-NC. Additionally, we identified an association between SVD in the temporal lobe and amygdala-predominant αSyn pathology, and highlighted its link with LATE-NC. Furthermore, our analysis showed that the *APOE* ε4 allele is associated with increased extracellular and capillary Aβ deposition but is not directly influenced by pTau burden or amygdala-predominant αSyn.

Consistent with previous studies,^4–6^ we found that mixed pathologies are highly prevalent in the brains of older adults. Only 32% of individuals with moderate to severe AD pathology had neither LATE-NC nor αSyn co-pathology. Individuals with AD and both co-pathologies showed significantly greater cognitive impairment than other patient groups, supporting previous findings that those with ‘quadruple’ lesions experience a more aggressive decline in cognitive function.^47^ Although pTau showed the strongest direct association with global cognitive decline, Aβ, CAA, and LATE-NC each contributed independent effects as well.

We observed that neuronal loss in CA1 subfield of hippocampus among older adults did not follow a binary pattern of intact tissue versus complete loss, i.e. hippocampal sclerosis, but instead occurred along a continuum. LATE-NC emerged as the strongest predictor of hippocampal damage, consistent with prior studies.^11,48–50^ However, tau also showed an independent effect in the full-cohort model, supporting findings from imaging studies that both pathologies contribute to MTL degeneration.^51^ Additionally, αSyn showed an independent association with CA1 degeneration—an effect we previously suspected but were unable to confirm in our smaller cohort.^16^

Using structural equation modeling, we found that the investigated proteinopathies were interconnected, consistent with evidence from cellular, animal, and in vivo studies suggesting that protein aggregates may promote misfolding and accumulation across pathologies.^14,15,17–19^ The strongest associations were identified between Aβ plaques and other lesions such as CAA, tau, and LATE-NC. We also confirmed a close relationship between LATE-NC and amygdala-predominant αSyn.^16^ Experimental studies have demonstrated that Aβ and αSyn can enhance tau seeding and aggregation,^14,15,52^ while αSyn has also been implicated in the deposition and seeding of TDP-43 pathology.^15,19^ Overall, Aβ and amygdala-predominant αSyn appear to play a less direct role in neuronal loss than tau and LATE-NC, but may play a crucial role in facilitating the aggregation and propagation of these lesions.

Next, we investigated the role of *APOE* ε4 in the network of neuropathologies. We confirmed a direct relationship between *APOE* ε4 and extracellular Aβ deposition, as well as a strong association with capillary CAA.^34,40,53^ In contrast, the general severity of CAA was not related to *APOE* ε4 but showed a modest association with cognitive impairment and was linked not only to Aβ, but also to pTau and amygdala-predominant αSyn. These findings suggest distinct underlying mechanisms for capillary and non-capillary CAA. Moreover, they support the hypothesis that one of CAA’s harmful effects is its role in exacerbating protein aggregation, for example, by stiffening blood vessels and restricting blood flow, thereby impairing perivascular clearance.^54,55^

We found no evidence of a direct link between *APOE* ε4 and Braak NFT staging. While mouse and cell models suggest that *APOE* ε4 independently promotes pTau accumulation,^56^ human studies align with our findings, showing no effect on pTau pathology after accounting for Aβ.^57–59^ However, since Braak NFT staging is a broad measure of pTau spread there is still a possibility that *APOE* ε4 is specifically associated with pTau burden in MTL regions, as suggested by some studies.^60^ Additionally, we found no relationship between *APOE* ε4 and amygdala-predominant αSyn, which contrasts with previous studies reporting that *APOE* ε4 exacerbates αSyn pathology independently of Aβ.^61,62^ This may be due to our more rigorous adjustment for Aβ lesions, or it could suggest that *APOE* ε4 primarily influences other variants of αSyn.

Finally, we cannot rule out a potential association between *APOE* ε4 and LATE-NC, as previously reported in the literature,^11,63^ given that this relationship in our cohort approached statistical significance. However, further research is needed to clarify the relationship between LATE-NC, *APOE* ε4, and Aβ, the latter of which showed an unexpectedly strong association with LATE-NC in our cohort. While previous studies have suggested that TDP-43 may inhibit Aβ fibrillization, resulting in the accumulation of toxic Aβ oligomers and increased neurotoxicity,^64^ this mechanism does not fully explain the observed LATE-NC association with overall Aβ burden.

We also systematically examined other pathologies that may contribute to global or hippocampal degeneration. Among cerebrovascular lesions, we found that SVD in the temporal cortex and atherosclerosis in the circle of Willis were both associated with CA1 neuron loss, aligning with previous findings.^23,28^ The CA1 region is particularly vulnerable to hypoxic damage,^65^ and cerebrovascular lesions may contribute to neuronal loss by reducing blood flow.^55^ However, in a comprehensive model, the association between SVD and CA1 damage appeared to be mediated by its links to LATE-NC and amygdala-predominant αSyn. While previous studies have established a connection between LATE-NC and SVD,^50,66,67^ the relationship between SVD and αSyn remains unclear and requires further investigation. Given that hypoxia has been shown to promote αSyn aggregation and enhance its toxicity,^68^ this link is biologically plausible. Moreover, SVD was associated with atherosclerosis and infarction, suggesting it may serve as a key mediator between cerebrovascular damage and the broader spectrum of AD neuropathology.

Interestingly, we found that Hirano bodies were positively associated with CA1 neuronal density, suggesting a potential protective role against hippocampal damage. These rod-shaped intracellular aggregates, primarily composed of actin,^69^ have been linked to neuropathological changes such as NFTs in AD.^70^ Consistent with our findings, studies using Hirano body models suggest that these inclusions may help reduce cell death associated with tau and the amyloid precursor protein intracellular domain.^71^ However, further research is needed to clarify their function in neurodegenerative processes.

In contrast, other lesions, including ARTAG, AGD, hippocampal vascular calcifications, and multiple old infarctions, showed no clear association with hippocampal or global brain degeneration, nor did they exhibit any discernible trends. Despite its established link to LATE-NC,^72^ ARTAG showed no indirect effects in our analysis. Instead, in our cohort, ARTAG was associated only with temporal SVD, not with LATE-NC. AGD was not associated with any measures of brain degeneration and, unexpectedly, was negatively correlated with AD-related lesions. This finding contributes to the ongoing debate on whether AGD represents a distinct neuropathological disease or if these tau lesions are largely benign.^73,74^

Neuronal density serves as a reliable marker of neurodegeneration and hippocampal atrophy, but manual counting is time-consuming and affected by variability in staining techniques, anatomical sampling, and cell morphology. While algorithms exist for neuron quantification from Nissl staining,^75^ this staining is incompatible with immunohistochemistry. To overcome this limitation, we developed a neuronal classifier for hematoxylin-stained slides. Its accuracy matches that of independent human raters, providing a reproducible and efficient approach for assessing pyramidal neuronal loss in large cohorts.

This study has several limitations. First, sample sizes varied across analyses. While core parameters were measured in all 480 cases, other measures (e.g., brain weight, CDR) were not available for the entire cohort. As a result, models based on incomplete samples may be underpowered and should be interpreted with caution. Second, the structural equation modeling assumed unidirectional pathways, which may oversimplify the complex biological interactions between protein aggregates. Reciprocal interactions, such as those between Aβ and pTau or between TDP-43 and pTau, exist but were not fully captured. Notably, the αSyn → LATE-NC pathway had to be reversed to achieve model fit, even though this reversal does not align with the suspected biological directionality. Finally, the parameters used to estimate tau, Aβ, and LATE-NC were based on lesion distribution (i.e., staging schemes) rather than (semi)quantitative assessments. This approach may influence the results, as lesion spread may not fully reflect the local pathological burden.

Overall, our findings indicate that AD pathology extends beyond Aβ and pTau, incorporating limbic-predominant variants of αSyn and TDP-43 pathologies, all of which contribute to brain degeneration. While pTau and LATE-NC were the strongest predictors of neuronal loss, Aβ and αSyn may play an additional critical role by promoting the aggregation and spread of other pathological proteins. These results underscore the need for improved in vivo detection of brain lesions to enhance patient stratification. They also highlight the importance of personalized treatment strategies that consider the complex interplay between protein aggregates, enabling more effective therapeutic interventions.

## Consent statement

All human donors in the study were fully informed and provided their consent to participate in accordance with ethical guidelines and the laws of Belgium, Germany, and the United Kingdom. All experiments were conducted following ethical clearance from the Ethics Committee Research UZ / KU Leuven (identifier: S65147).

## Supporting information

This study was supported by grants from the Fond Wetenschappelijk Onderzoek (FWO, Belgium): G0F8516N (DRT), G065721N, and G024925N (DRT, KS); KU Leuven Onderzoeksraad (Belgium): C14/17/107 (DRT), C14/22/132 (DRT), and C3/20/057 (DRT); Deutsche Forschungsgemeinschaft (DFG, Germany): TH-624-4-1, TH-624-4-2, TH-624-6-1 (DRT); Alzheimer Forschung Initiative (AFI, Germany): #10810, #13803 (DRT); and the Alzheimer’s Association (USA): 22-AAIIA-963171 (DRT). SOT was supported by KU Leuven Internal Funding (PDMT2/21/069), the BrightFocus Foundation (A2022019F), Fonds Wetenschappelijk Onderzoek (1225725N), and the Alzheimer’s Association (AARF-24-1300693). MO received support from the EU Joint Programme-Neurodegenerative Diseases network Genfi-Prox (01ED2008A), the German Federal Ministry of Education and Research (FTLDc 01GI1007A), the EU Moodmarker program (01EW2008), the German Research Foundation (DFG) (SFB1279), the Foundation of the State of Baden-Württemberg (D.3830), Boehringer Ingelheim Ulm University BioCenter (D.5009), the Thierry Latran Foundation (D.2468), and the Roux Program of the Martin Luther University Halle-Wittenberg.

## Acknowledgements

The authors thank GE Healthcare for providing samples for this study and Petra Weckx for technical assistance. We appreciate Grzegorz Walkiewicz and Bas Moonen for their feedback on the content and figures. We also acknowledge Kamil Sekut for proposing the idea behind the machine-learning algorithm for neuron detection. Finally, we thank the patients and their families for their contributions.

