## Supplementary Figures for "Alzheimer’s disease and its co-pathologies: implications for hippocampal degeneration, cognitive decline, and the role of *APOE* ε4"

**List of content**


**Figure 1.** Example results of the automated detection and classification algorithm applied to posterior CA1 region images.
**Figure 2.** Scatter plot showing neuron detection counts by the algorithm and an independent rater.
**Figure 3.** Distributions of neuronal counts and brain weight, stratified by cognitive impairment.
**Figure 4.** Heatmaps displaying the results of Series 1 and 2 of linear regression analyses, including statistical significance and the number of cases used.
**Figure 5.** Heatmaps showing exact standardized coefficients from two series of multiple linear regression analyses.
**Figure 6.** Results of supplementary structural equation models.

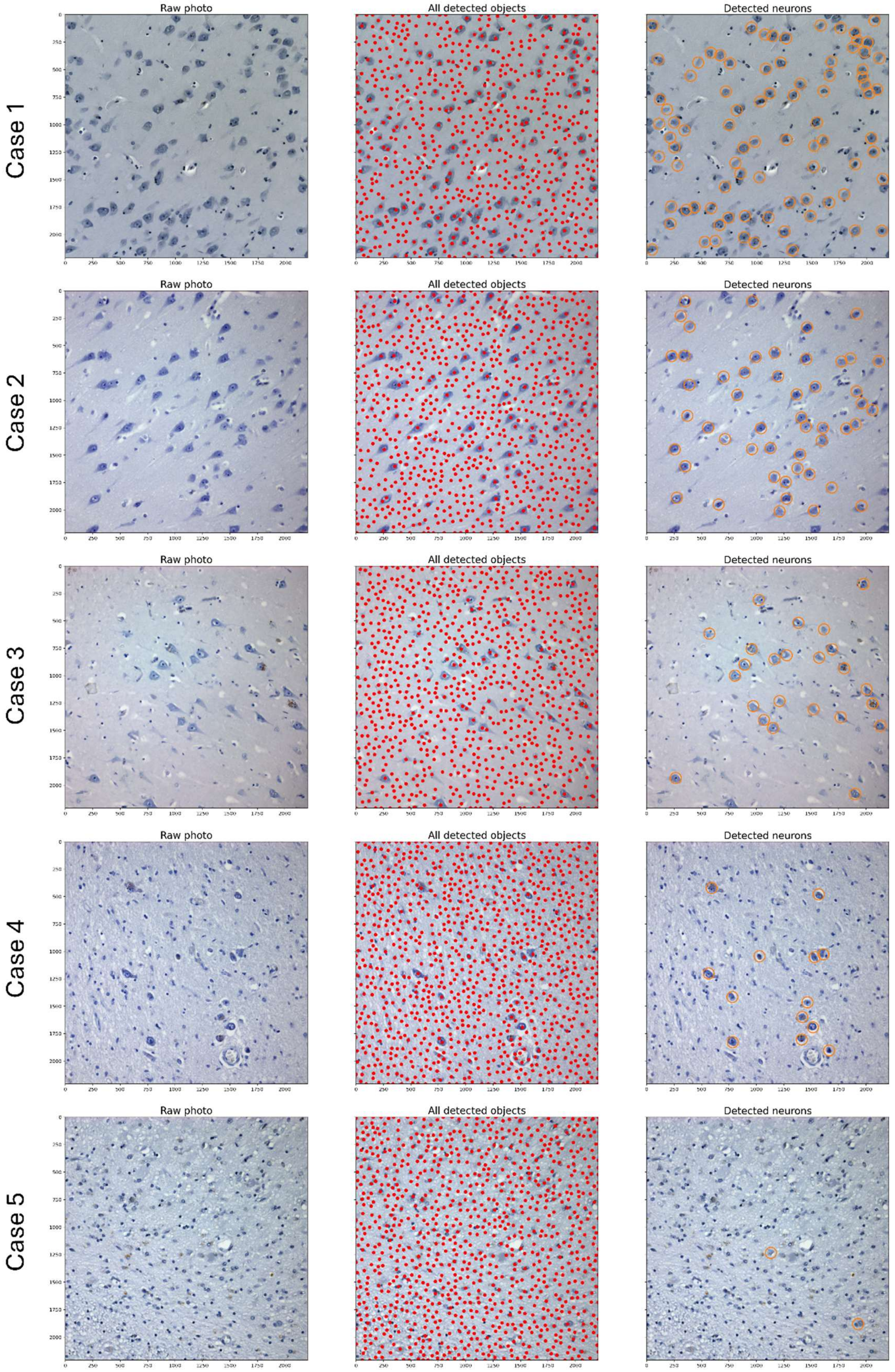
Raw photo
All detected objects
Detected neurons
Case 1
Case 2
Case 3
Case 4
Case 5

**SUPPLEMENTARY FIGURE 1**

Example results of the automated detection and classification algorithm applied to posterior CA1 region images at x200 magnification. **Left panel**: Raw microscopy images showing cellular structures within the CA1 region. **Middle panel**: Detected objects, including neurons and non-neuronal structures, as identified by the automated detection algorithm. Each object was subsequently classified using a trained neural network into "neuron" or "non-neuron" categories. **Right panel**: Objects identified and classified as neurons by the algorithm. The five cases displayed correspond to decreasing CA1 neuronal densities per $mm^2$: 301.2, 202.7, 141.3, 41.1, and 15.9, reflecting a gradient of increasing neuronal loss across the samples.

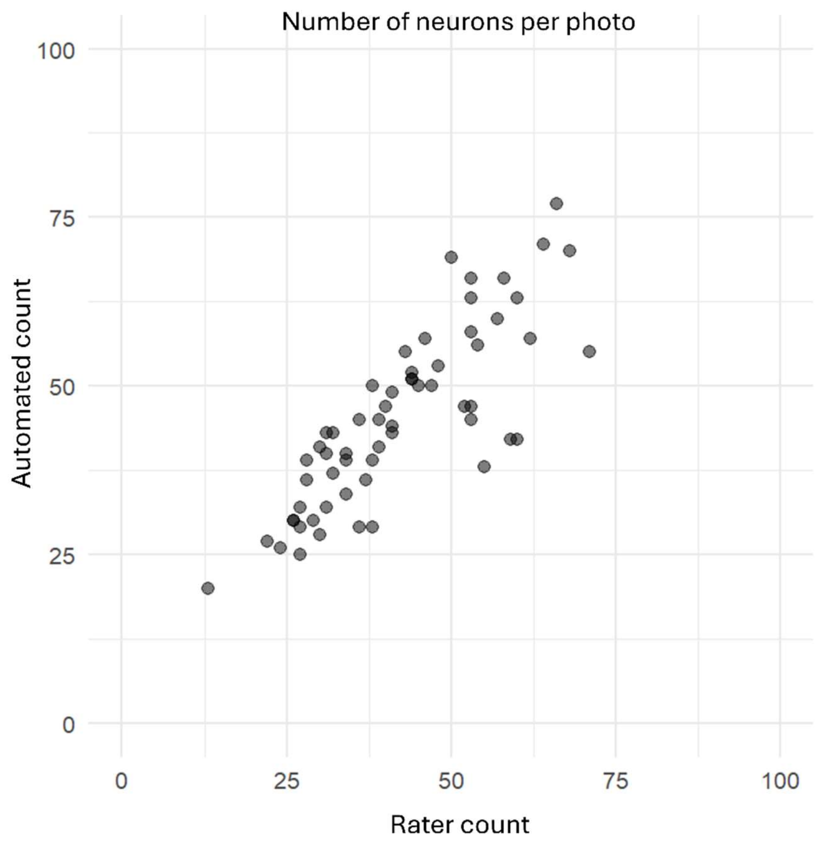


**SUPPLEMENTARY FIGURE 2**

Scatter plot illustrating a monotonically increasing relationship between neuronal counts in 60 tested images as detected by the automated detection algorithm and those assessed by an independent human rater. The Spearman's rho for this correlation was 0.83.

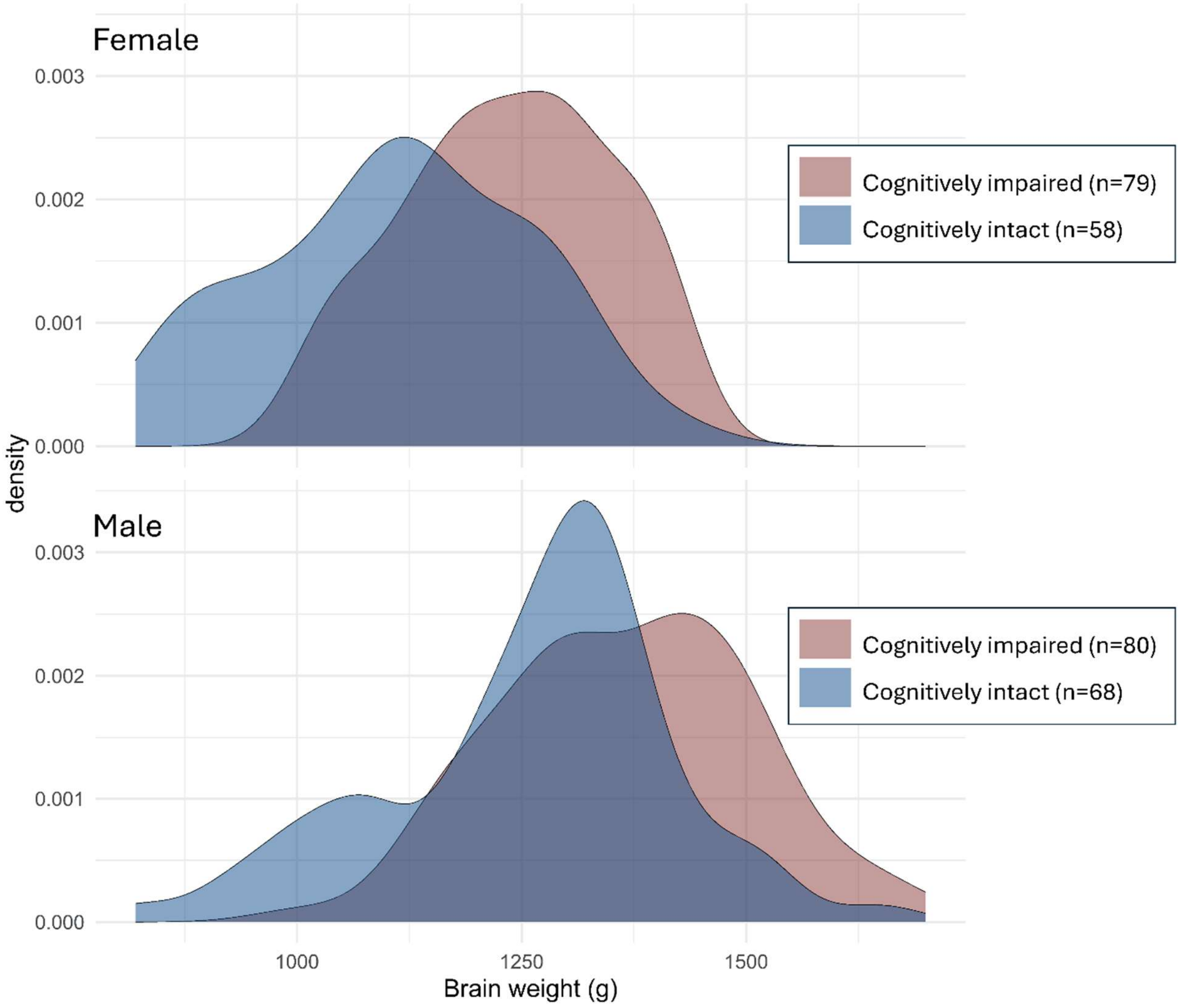


**SUPPLEMENTARY FIGURE 3**

Density plot showing sex-stratified distributions of brain weight (grams) for cognitively impaired and cognitively intact individuals. The remaining cases (n = 195) lacked data on cognitive impairment status or brain weight.

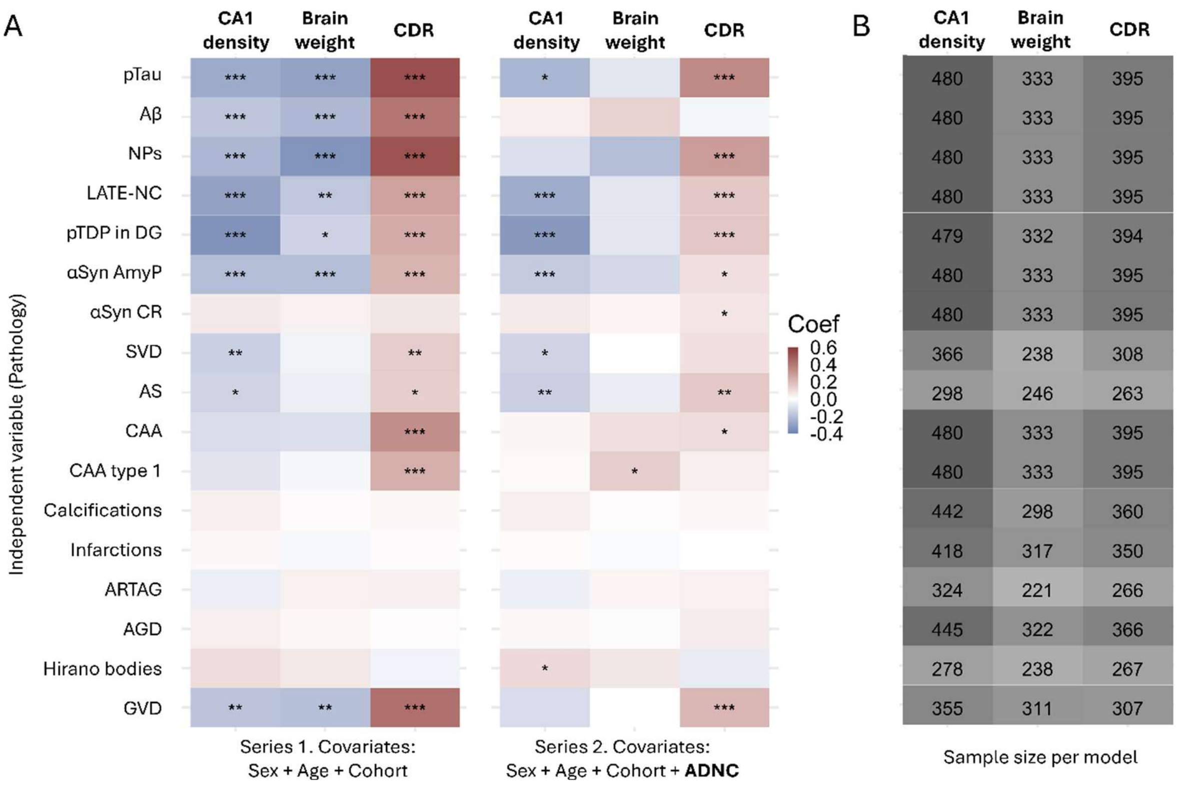


**SUPPLEMENTARY FIGURE 4**

**A.** Heatmap displaying standardized coefficients from multiple linear regression analyses, with brain damage measures as dependent variables and neuropathological lesions as independent variables. The first series of regressions was adjusted for age at death, sex, and cohort. The second series included additional adjustment for the overall severity of Alzheimer's disease pathology. Asterisks on the tiles denote p-values after Benjamini-Hochberg correction within each panel: $*p < 0.05$, $**p < 0.01$, $***p < 0.001$. **B.** Number of cases included in each model.

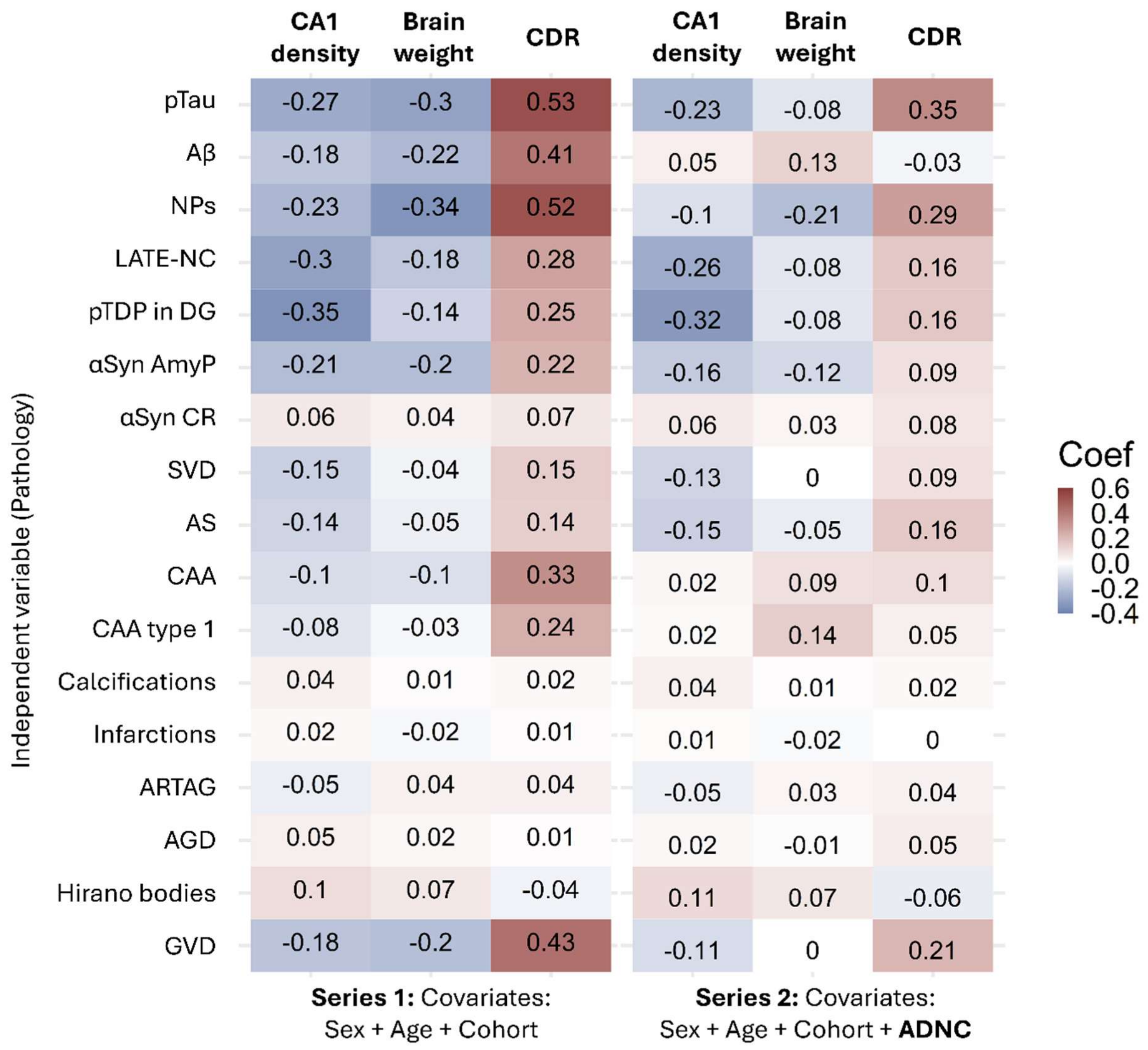


**SUPPLEMENTARY FIGURE 5**

Heatmap with exact values of standardized coefficients from multiple linear regression analyses, with brain damage measures as dependent variables and neuropathological lesions as independent variables.

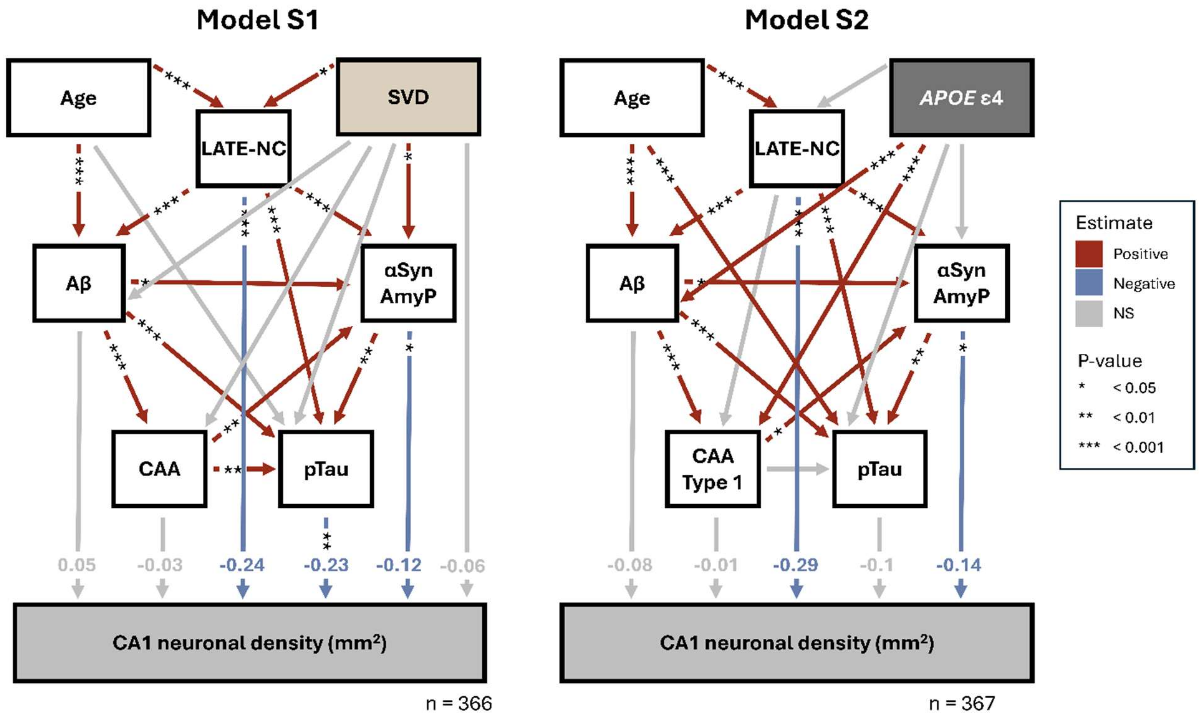


**SUPPLEMENTARY FIGURE 6**

Results of supplementary structural equation models. Each modeled directional relationship is represented by an arrow on the graph. The color of the arrow indicates the direction of the obtained coefficient in the analysis, with statistically non-significant coefficients shown in gray. The standardized estimate for paths involving brain damage parameters is displayed at the end of each arrow.
